# Savanna chimpanzees (*Pan troglodytes verus*) in Senegal react to deadly snakes and other reptiles: Testing the snake detection hypothesis

**DOI:** 10.1101/2022.09.04.506548

**Authors:** Jill D. Pruetz, Thomas C. LaDuke, K. Dobson

## Abstract

The hypothesis that dangerous snakes served as evolutionary selective pressures on traits characterizing the Order Primates (Snake Detection Hypothesis, SDH), specifically vision and aspects of the brain, has received recent attention. We provide data on 52 encounters between chimpanzees (*Pan troglodytes verus*) in a savanna landscape and snakes and other large reptiles at the Fongoli site in Senegal, over the course of eight years. These encounters yielded data on 178 interactions between identified individual chimpanzees and reptiles. The majority of encounters with identified reptiles (62%) involved potentially deadly snakes. Constrictors of the family Pythonidae were encountered more frequently than any other reptile. Chimpanzees exhibited a variety of reactions to reptiles, ranging from attacking with objects to ignoring them. Even reptiles other than snakes were met with some degree of alarm when they were in water or moving quickly. We assessed variables that may affect apes’ reactions, as well as the age-sex class of chimpanzees. As a test of Isbell’s snake detection hypothesis, we examined chimpanzees’ reaction intensity to venomous versus constricting snakes but found no difference. We did find significant age-sex differences in chimpanzees’ reactions to snakes, with adolescent males exhibiting higher-intensity reactions than adult males and females and adolescent female chimpanzees. Chimpanzees reacted at significantly higher intensities when snakes were arboreal in location, but reptile species, size, and activity did not significantly influence individuals’ reactions to snakes. We also report one inferred encounter between an adult female chimpanzee and a venomous snake, which led to her death. Our data suggest that snakes are significant threats to chimpanzees in savanna environments and support the hypothesis that danger from snakes could have exerted significant evolutionary pressure on the Order Primates.

## Introduction

The Snake Detection Hypothesis (SDH) posits that dangerous snakes were influential in the evolution of the visual system and fear module in the brain of primates [1–2] and prompts closer study of living primates’ interactions with snakes. Isbell [1–2] suggests that characters of the anthropoid primates in particular are related to the emergence of venomous snakes but points out that there are little data available to distinguish between the reactions of primates to constricting versus venomous snakes (Table 1). Of such published accounts, primates were able to escape constrictors almost as often as not, but more lethal events were associated with constrictors than venomous snakes (Table 1). However, encounters with constrictors are more likely to be observed because of the size of snake and the duration of the encounter compared to primates’ encounters with venomous snakes. Additionally, the sample size across the Order Primates is sparse, and most published reports contain only events in which individuals were killed or attacked by snakes [3–24]. Fewer still report primates’ general reactions to snakes, save for captive or experimental studies, where considerable research indicates that at least some primate species are “programmed” to fear snake-like objects, albeit with some degree of learned modification of behavior possible [25–28]. Conversely, a study of orbital convergence in primate species at risk of predation by venomous versus constricting snakes failed to support the SDH [29].

**Table 1.** Published accounts of encounters between non-human primates and snakes.

| **Primate species** | **Snake species** | **Snake type\*** | **Outcome** | **Reference** |
| --- | --- | --- | --- | --- |
| *Nycticebus coucang* | *Python reticulatus* | C | Lethal | [3] |
| *Microcebus murinus* | *Sanzinia madagascariensis* | C | Non-lethal; physical conspecific aid | [4] |
| *Propithecus coquereli* | *Acrantophis madagascariensis* | C | Non-lethal; physical conspecific aid | [5] |
| *Tarsius spectrum* | *P. reticulatus* | C | Lethal; non-physical conspecific aid | [6] |
| *Callithrix jacchus* | *Bothrops leucurus* | V | Lethal | [7] |
| *C. aurita* | *Bothrops jaracara* | V | Lethal | [8] |
| *C. pencillata* | *Boa constrictor* | C | Lethal (2 juveniles); 2 adults attack snake | [9] |
| *Leontopithecus rosalia* | *Boa constrictor* | C | Non-lethal | [10] |
| *Saguinus mystax* | *Boa constrictor* | C | Non-lethal; physical conspecific aid | [11] |
| *S. mystax* | *Eunectes murinus* | C | Lethal | [12] |
| *S. fuscicollis* | *Philodryas nattereri* | V | Non-lethal | [13] |
| *Callicebus discolor* | *Boa constrictor* | C | Non-lethal | [14] |
| *Alouatta puruensis* | *Boa constrictor* | C | Lethal | [15] |
| *Alouatta palliata* | *Boa constrictor* | C | Non-lethal | Fedigan, cited in [16] |
| *Chiropotes utahickae* | *Boa constrictor* | C | Lethal | [17] |
| *Cebus capucinus* | *Boa constrictor* | C | Non-lethal & lethal; non-physical conspecific aid in the latter | [18] |
| *Cebus capucinus* | *Boa constrictor* | C | Non-lethal; physical conspecific aid | [19] |
| *Sapajus* | *Corallus enydris* | C | Non-lethal | [20] |
| *Saimiri sciureus* | *Boa* | C | Lethal | [21] |
| *Cercopithecus mitis stuhlmanni* | *Bitis gabonica* | V | Lethal | [22] |
| *Cercopithecus mitis albogularis* | *Dendroaspis polylepsis* | V | Lethal, inferred | [23] |
| *Chlorocebus aethiops* | *Python* | C | Lethal | [24] |
| *Macaca nemestrina* | *P. moluras* | C | Lethal | [25] |
| *Trachypithecus auratus* | *Python reticulatus* | C | Non-lethal, encounter | [26] |
| *Pan troglodytes verus* | *Dendroaspis polylepsis* | V | Lethal, inferred | This study. |

\*constricting = C, venomous = V

Even though relatively few snake species pose a significant predator threat to primates, many more species nonetheless pose a lethal threat due to the effects of their venom. Isbell’s [1] hypothesis to explain the evolution of primates and, specifically, their well-developed visual system and fear module therefore is informed in part on the few naturalistic observations of primates’ predators and their potential prey thus far reported. We provide data on chimpanzees’ reactions to snakes and other reptiles in a savanna-woodland habitat at the Fongoli, Senegal study site. We use data on individual chimpanzees of various age and sex classes as they encounter and react to reptiles of different types, location, activities and size to explore patterns that can inform our understanding of the degree to which chimpanzees regard potentially dangerous snakes as threats. Specifically, we test the hypothesis that chimpanzees will react differentially to venomous versus constricting snakes. Our data can help contribute to a growing body of work on the subject, set in motion by Isbell’s [1] hypothesis.

## Methods

### Study site and subjects

The Fongoli chimpanzee community ranges within the Kedougou (formerly Tambacounda) Department in southeastern Senegal (12º40N, 12º13W). The savanna environment here is a mosaic of woodland, grassland, bamboo and gallery forest habitats [30]. The extensive dry season lasts over seven months, and rainfall usually averages less than 1000 mm annually [31]. Maximum temperature in the dry season exceeds 40°C. The Fongoli chimpanzee community varied annually between 29 and 36 individuals during the study period, from June 2005 to November 2015 (10.25 years), averaging 32.3 individuals during this time. Most community members were habituated in 2005, but some adult females were semi-habituated, in that they exhibited signs of nervousness around observers when adult males were absent. Age-sex classes used for analyses were as follows: infant males and females <4 years; juvenile males and females >4 years, <7 years; adolescent males >7 and <15 years; adolescent females >7 years until giving birth; adult males >15 years; adult females, following first infant birth.

Relatively few natural predators remain at Fongoli, where chimpanzees are sympatric with humans who practice horticulture, hunting and some gathering. Humans are usually treated to some degree as a threat, and chimpanzees’ reactions usually range from hiding to silent fleeing. Predators that have been exterminated or otherwise are no longer found at Fongoli include lion (*Panthera leo*), cheetah (*Acinonyx jubatus*) and wild dog (*Lycaon pictus*), but older chimpanzees most likely experienced these predators in previous decades (M. Camara, pers. comm.). Rare evidence, including a witnessed encounter between chimpanzees using weapons against a leopard (*Panthera pardus*) as well as camera trap data and one sighting, indicates that this big cat is resident in the area (Pruetz & Boyer, in prep.). Spotted hyenas (*Crocuta crocuta*) are seen periodically, and treated as predators by chimpanzees (i.e., mobbed; Pruetz & Boyer, in prep.). A hippopotamus (*Hippopotamus amphibius*) in the Gambia River has also been treated as a predator as it surfaced to breathe (Fongoli Savanna Chimpanzee Project, unpub. data). Crocodiles (*Crocodilus niloticus*) are present in the Gambia River and, seasonally, in its tributaries, but the individuals of this species that enter the tributaries within the chimpanzee’s home range are apparently small (generally less than one meter in length) based on hunting (by human) traces identified by TCL and observations by FSCP personnel.

While only one reptile species poses a significant predator threat to Fongoli apes (i.e., the rock python, *Python sebae* and perhaps *P. regius*), a number of snakes produce venom that poses a lethal threat to even adult chimpanzees, i.e., mamba (*Dendropaspis polylepis*), spitting cobra (*Naja nigricollis* or *katiensis*), black cobra (*Naja melanoleuca*), spotted night adder (*Causus maculatus*), puff adder (*Bitis arietans*), and saw-scaled viper (*Echis ocellatus*). The elapid snakes can be categorized as actively foraging species and are generally mobile when encountered during daylight hours when chimps are likely to encounter them. The vipers and pythons are largely ambush predators and are generally coiled and immobile when chimps encounter them.

### Data collection

To provide some indication of the frequency with which human observers encounter various species of snakes at Fongoli, we provide a summary of encounters between observers and venomous snakes both within and outside of those that occur during follows of chimpanzees. Sightings are stochastic, and the frequency depends on the number of observers on site. However, on average, only one to two observers in addition to JDP are at Fongoli at any one time. Most identification of snakes and other herpetofauna was made by JDP in conjunction with TCL, a herpetologist. Such identification included discussions of specific traits observed, such as general morphology and color patterns and, in the case of the black mamba and spotted night adder (*Causus maculatus*), when both JDP and TCL were on site. Some snakes were identified from photographs. Most data stem from JDP’s records, which account for approximately 66 months (~1850 days) spent in the field during the study period: June 2005-November 2015. The study period began following habituation of adult males to nest-to-nest follows from a distance of <20m in early 2005. Snake observations are biased towards early (dusk) and late evening and early morning (pre-dawn) time periods, when researchers were traveling to or from chimpanzee nesting sites. Researchers rarely saw snakes that chimpanzees did not first detect during chimpanzee follows, and the former usually involved moving reptiles. We recorded encounters between chimpanzees and other larger reptiles as well. We defined a large reptile as any snake or other reptile larger than the common Agama (*Agama agama*) lizard, which is ubiquitous but which apes were never seen to react to save for one instance in which an adolescent male attempted to capture one but failed.

When chimpanzees were observed to encounter large reptiles, the habitat type, type of snake, individual chimpanzees involved and their reactions, as well as the location and behavior of the snake were recorded (S1, Tables 1 & 2). Encounters were scored according to one of the following categories, listed from low to high-intensity in terms of reaction to the reptile:

1. Ignore – chimpanzee is apparently aware of reptile but exhibits no signs of fear or is curious, in that it appears to see reptile but exhibits no signs of fear or anxiety;
2. Vigilant peering – chimpanzee looks alertly at reptile, may exhibit piloerection, includes approaching reptile to peer; corresponds to [32] ‘gaze toward’;
3. Vocalize – chimpanzee emits (a) alarm calls (“waaa”) or (b) inquisitive calls (“huu”) upon detecting reptile [33; and see Supplemental Video], includes approaching to vocalize; corresponds to [32] ‘alarm call’;
4. Avoid – chimpanzee moves away from reptile or leaves the area, indicative of avoiding possible interaction with the reptile;
5. Mob – chimpanzee threatens reptile, including (a) display at, (b) hits at with object or throws object at reptile, or (c) strikes reptile with object; corresponds to [7] ‘mob’.

**Table 2.** Reptile species identified at Fongoli and relative estimates of abundance based on encounter rate (includes encounters when observers were with chimpanzees, traveling to and from chimpanzee nest sites, and in Fongoli village).

| Species common name | Scientific name | Relative frequency of sightings by observers | Relative density in region following [34]* |
| --- | --- | --- | --- |
| ***Venomous*** |  |  |  |
| Black mamba | *Dendroaspis polylepis* | Yearly | Rare |
| Black-necked spitting cobra | *Naja nigricollis* | Yearly, usually at camp | Rare |
| Black or forest cobra | *Naja melanoleuca* | < 1 per year in most years | Rare |
| Puff adder | *Bitis arietans* | Multiple times yearly | Occasional to frequent |
| Spotted night adder | *Causus maculatus* | Multiple times yearly likely | Occasional |
| Saw-scaled viper* | *Echis ocellatus* | Multiple times yearly likely | Occasional |
| Twig snake | *Thelotornis kirtlandi* | < 1 per year | Rare |
| Boomslang | *Dispholidus typus* | < 1 per year but I.D. difficult | Rare |
| **Non-venomous** |  |  |  |
| Rock python | *Python sebae* | >1 per year but varies annually | Rare |
| Ball python | *Python regius* | Rare | Rare |
| House snake | *Lamprophis* sp. | Usually yearly | Occasional |
| Sand snake** | *Psammophis* sp. | Multiple times yearly | Occasional |
| Rufous-beaked snake | *Rhabdophis* | Rare (once) | Rare |
| Shovel-snout snake | *Prosymna* | Multiple times yearly | Occasional |
| Thread snake | *Leptotyphlops* sp. | Less than once per year (in well) | Rare |
| Green or bush snake | *Philothamnus* | Rare to multiple times per year but varies annually | Rare |
| Nile monitor lizard | *Varanus niloticus* | Common | Frequent |
| Savanna monitor lizard | *Varanus exanthematicus* | Less common than savanna, I.D. not always positive between species | Rare to frequent |
| Nile crocodile | *Crocodylus niloticus* | Rare | Rare |
| Agama lizard | *Agama agama* | Ubiquitous (usually multiple times daily) | Very frequent |
| African spurred tortoise | *Geochelone sulcata* | Rare (less than twice) | Rare |
| Soft-shelled turtle | *Cyclanorbis senegalensis* | Rare (once) | Rare |
| Hinge-backed tortoise | *Kinixys homeana* | Rare | Rare |
| Unidentified turtle | ? | Usually yearly | Occasional |

\***Rare** = never observed or <1 per 6 research months. **Occasional** = once per 1-6 research months. **Frequent** = once per month. **Very frequent** = >10 observations per month (Hockings et al., 2012 following Ross et al. 2009).

Often, observers were alerted to the presence of a snake by the chimpanzees’ calls. Snakes elicit an acoustically specific alarm-call from the Fongoli chimpanzees just as they do from many other primates [33; Pruetz, personal observation]. Such an alarm-call is distinguishable by even relatively naive observers, although no experimental verification of this predator-specific vocalization has been conducted. Researchers investigated “inquisitive huu” vocalizations for possible snake interactions less extensively during early years of the project compared to later years. In cases where the ‘huu’ vocalization escalated into ‘waaa’ alarm calls or when multiple individuals emitted such vocalizations, observers investigated the context in all years. However, there are likely more potential snake-chimpanzee interactions occurring than observers recorded, especially during the early years of the study. In 2015, snake alarms along with behavior consistent with reactions to pythons were observed, but the observer was unable to detect a snake in the thick vine tangles where chimpanzees directed their aggression. In one of these cases, chimpanzees detected a medium-sized python in the same tree as a few days earlier, which the observer was also able to see. Similarly, several times the alarm call of individuals elicited the approach and vigilant behavior of other chimpanzees, but the observer was never able to verify the presence of a snake. These cases were not included in analyses.

### Data summary and analyses

For certain analyses, we lumped reaction types regarding intensity. We considered ‘peer’ and ‘ignore’ the lowest reaction intensity; a medium level of reaction included leaving, avoiding and vocalizing; the most intense reaction category included hit, hit at, charge and display.

We conducted regression analyses in SYSTAT to examine the effects of independent variables such as snake type (venomous, constricting), snake activity (mobile, immobile), and snake substrate (arboreal, terrestrial) on the intensity of chimpanzees’ reactions. We excluded male and female juvenile and infant chimpanzees for analysis because of small sample sizes and the assumption that, for these dependent offspring, their behavior would likely reflect the behavior of their mother. Only pythons were included in the constricting category, while black mambas, cobras and all vipers were included in the venomous category. The back-fanged sand snake was not included in the venomous category given the relatively low chance it would prove deadly to chimpanzees. The dependent variable examined was chimpanzee reaction type.

We conducted multinomial logistic regression analyses to examine the effects that venomous (cobras, mambas, unidentified elapids, vipers) versus constricting (python) snakes as well as chimpanzee age-sex class had on reactions to snakes. We used three categories of reactions to snakes (high [attack or charge], medium [vocalize at or avoid/leave] and low [vigilant peer or ignore] intensity) and classified individual chimpanzees as adult male, adolescent male, or female. We combined the adolescent female category with the adult female category since the adolescent female sample size was relatively small (see Figures 1 & 2) and was characterized by an empty cell (no medium-intensity interactions with constricting snakes). We also examined the relative significance of variables such as snake size (small [vipers other than puff adders, non-venomous snakes], medium [puff adders, mambas <2m in length] or large [pythons, mambas >2m, puff adders >1m]), location (arboreal or terrestrial), and activity (moving, immobile) on chimpanzee reactions.

**Figure 1.**
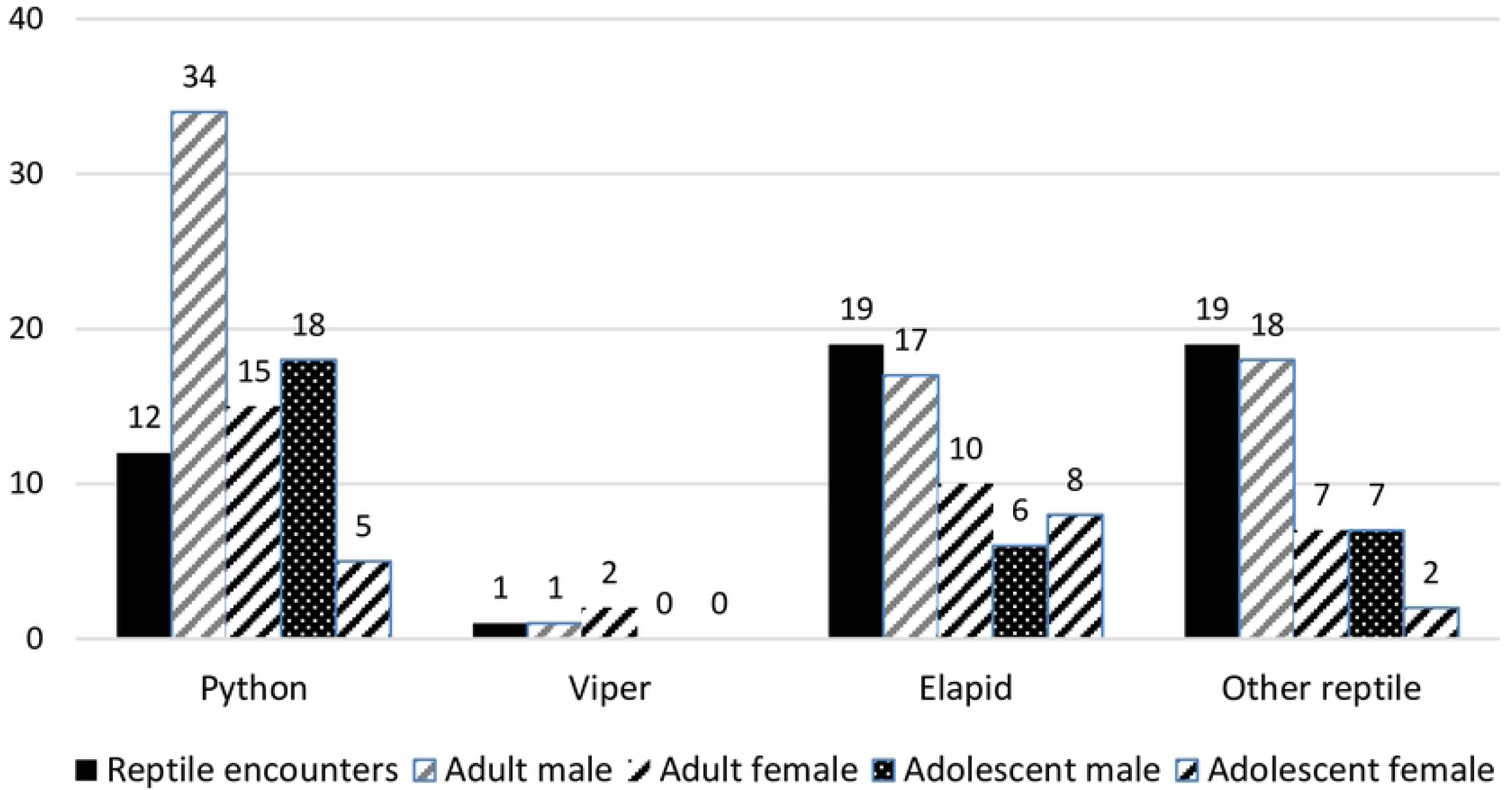
Frequencies of interactions between age-sex classes of chimpanzees in encounters with different types of large reptiles.

**Figure 2.**
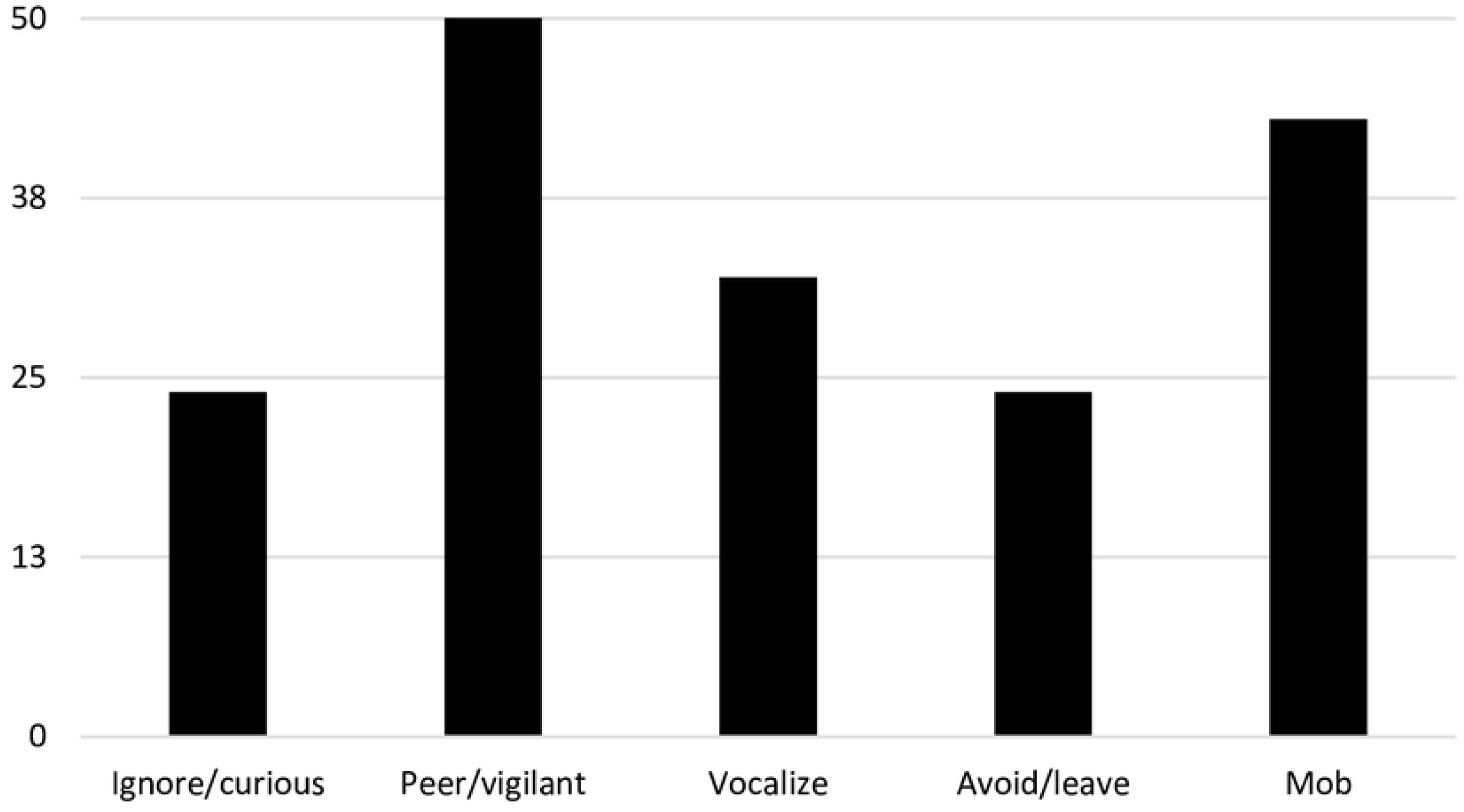
Frequency of the different types of reactions exhibited by individual chimpanzees towards large reptiles at Fongoli.

We conducted a binary logistic regression analysis to consider the effects of more variables on chimpanzees’ reactions to snakes. We used two categories of reactions to snakes (high [attack or charge, avoid/leave, or vocalize] and low [vigilant watch or ignore]) but four levels of age-sex class: adult male, adult female, adolescent male, adolescent female. We also examined the effects that snake type, location, size and activity had on overall chimpanzee reaction (i.e., age-sex classes lumped for analyses).

## Results

### Snake presence at Fongoli

Table 2 provides a summary of reptiles encountered at Fongoli, their relative observed frequency and comparison with other sites where similar data are available. The only snakes identified to species were pythons (*n*=14 cases), puff adders (*n*=10 cases involving 11 puff adders), cobras (*n*=2 cases), black mambas (*n*=5 cases), saw-scaled viper (*n*=1 case), and the striped sand snake (n=1 case). Tentative identifications were obtained for puff adders (*n*=1), an elapid (*n*=1) and a monitor lizard (*n*=1). An additional eight cases included unidentified snakes, although one of these was not any of the venomous species known in the area (unless it was an unrecognized back-fanged colubrid species), nor was it a python. Not included in Table 2 are several other cases of interest:

1. An interaction between a research assistant (D. Kante) and a venomous snake in the nearby (~10km) town of Kedougou. He was envenomated in the eye by a spitting cobra as he passed below a low-hanging branch. Immediate medical care resulted in rapid recovery.
2. A woman from a nearby village was collecting *Saba* fruit in the chimpanzees’ home range in July 2002 and was bitten by a snake while climbing in a tree crown. She was subsequently taken to the hospital in Kedougou. No other information on her condition was given, but it is assumed she survived or word would have reached project personnel.
3. Research assistants D. Kante and M. Sadiakho report one woman from their home village (Thiobo: within several kilometers of the study site) dying from a bite of what appeared to have been a saw-scaled viper.
4. Researcher M. Sadiakho reports that, in previous years, a large rock python hunted domestic dogs in the vicinity of his home village, Tenkoto, which lies just outside (~1km) of the home range boundaries of the Fongoli chimpanzees’ home range. Villagers ultimately killed the snake.
5. Researchers found one dead, mummified rock python more than three meters long that was tied to a tree within the Fongoli chimpanzees’ home range. This is a common means of dealing with snakes after they have been killed by local people (M. Sadiakho, pers. comm.).
6. After a juvenile male emited inquisitive ‘huuu’ vocalizations at shed snake skin, approximately one meter in length. Several other individuals peered at shed but continued. A young adolescent male examined then played with the shed skin for several minutes, also rubbing it over himself.
7. In 2019, Researcher M. Sadiakho was bitten by a small viper in a recently harvested peanut field in his home village of Tenkoto, near the edge of the Fongoli chimpanzee home range. He consumed tobacco upon the advice of local people, became nauseous and was taken to the Kedougou hospital, approximately 20 km away, where he was given antivenom. He recovered and was back at work after three days.

### Interactions between chimpanzees and reptiles

Chimpanzees at Fongoli appeared attracted to large reptiles, even when they were not the first individual to see them (S1, Table 2). Adult males most often interacted with large reptiles, although we were not certain that we recorded all instances of interactions and therefore did not analyze frequency of interaction according to age-sex class (Fig. 1). Adult males interacted with large reptiles 74 times, while adult females interacted with reptiles 36 times (Fig. 1). Adolescent males interacted with large reptiles 36 times, while adolescent females interacted with large reptiles 14 times (Fig. 1).

Chimpanzee reactions varied according to reptile species, though dangerous (venomous or constricting snakes) species were almost always met with some type of fearful reaction (S1, Tables 1 & 2). Other reptiles also elicited fear, especially when they startled chimpanzees, except for small lizards. Chimpanzees frequently encounter *Agama* lizards, which are ignored except for mild curiosity in one case and a failed attempt to capture one in another, even when these lizards exhibit display head-bobbing behaviors in response to chimpanzee presence or loud calls. Monitor lizards (*Varanus* sp.) were usually ignored, except for instances in which they startled chimpanzees or shared the same tree crown with them. Turtles (Testudines) were met with fear (piloerection and avoidance behavior) and aggressive behavior, especially when in water. A chameleon (*Chamaeleo* sp.) was treated cautiously although immature individuals attacked it, throwing and poking it.

Overall, the reactions of individual chimpanzee varied from the mildest of ignoring reptiles (*n*=18) to the most intense of physically attacking them with objects (*n*=21), a form of mobbing (Fig. 2). Vigilant peering (*n*=54) accounted for the greatest number of reactions, followed by mobbing (n=43 total mobbing events), vocal threat (*n*=35), avoiding or leaving the area (*n*=24) and non-contact aggressive behavior directed at the reptile (displaying or hitting at, *n*=15.

Of 52 encounters between apes and large reptiles at Fongoli, 40 were with snakes. Approximately 61% (*n*=32) of the encounters with identified snakes involved dangerous species, such as pythons or venomous snakes. One case included a back-fanged species (*Psammophis*), and the remaining six cases included unidentified snakes. Other reptiles encountered included monitor lizards, turtles, tortoises, and chameleons.

### Factors effecting chimpanzees’ reactions to dangerous snakes

Multinomial logistic regression analyses of individual chimpanzees’ reactions (high, medium, and low intensity) according to three chimpanzee age-sex classes (adult male, female [adolescent and adult], adolescent male) and dangerous snake type (venomous or constricting) revealed no significance in the overall model (*n*=114, X^2^=7.086, df=4, p=0.131) (Fig. 3). Including snake location (arboreal or terrestrial) showed that this variable affected chimpanzees’ reactions (95% CI=0.137-2.044, p=0.025, z-score = 2.242), but the model only approached significance levels (*n*=108, X^2^=10.718, df=6, p=0.097).

**Figure 3.**
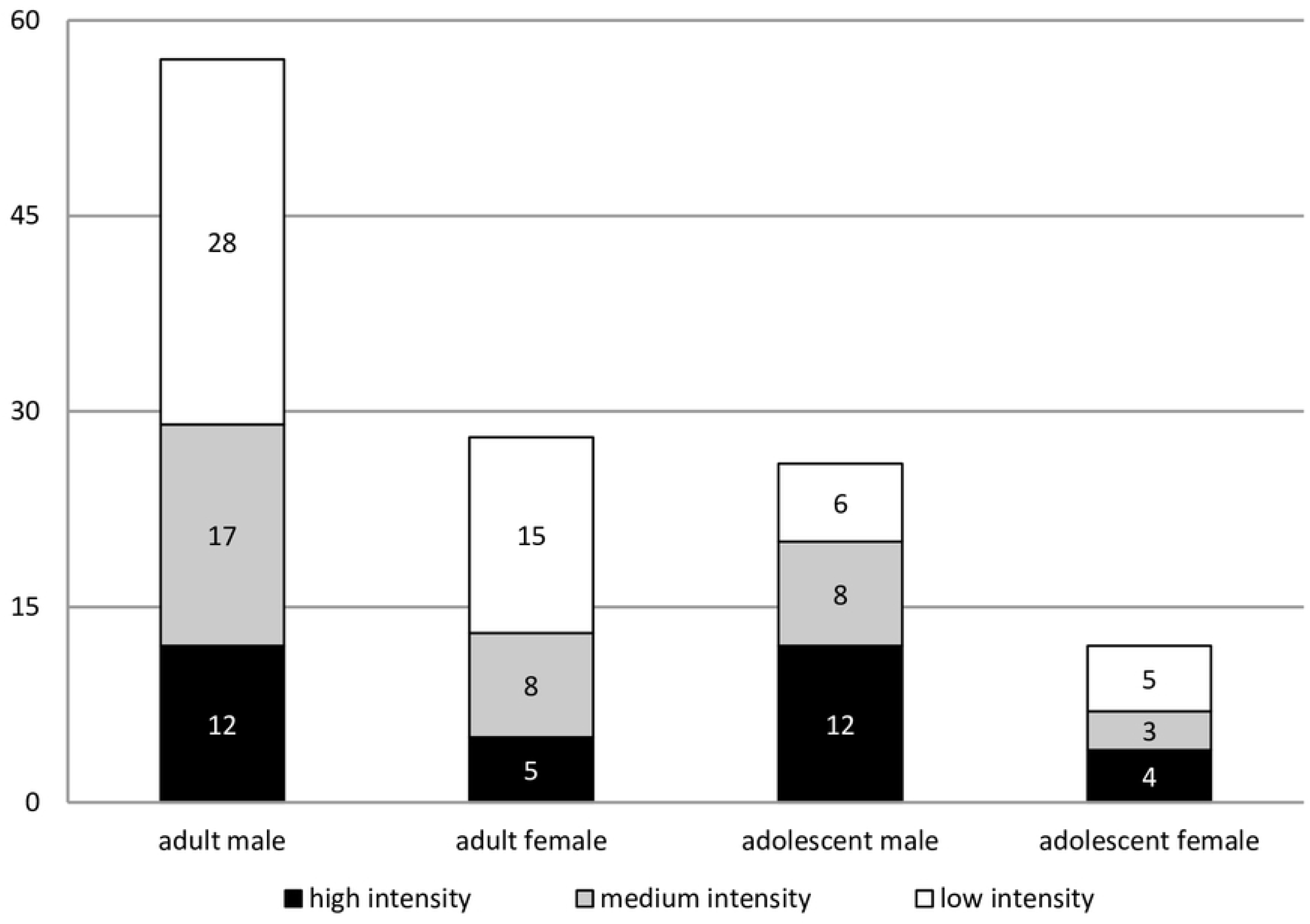
Frequency of reactions of various intensities by chimpanzees in different age-sex classes. *high intensity = hit, hit at, charge or display; medium intensity = leave, avoid or vocalize at; low intensity = peer, vigilant at or ignore.

**Figure 4.**
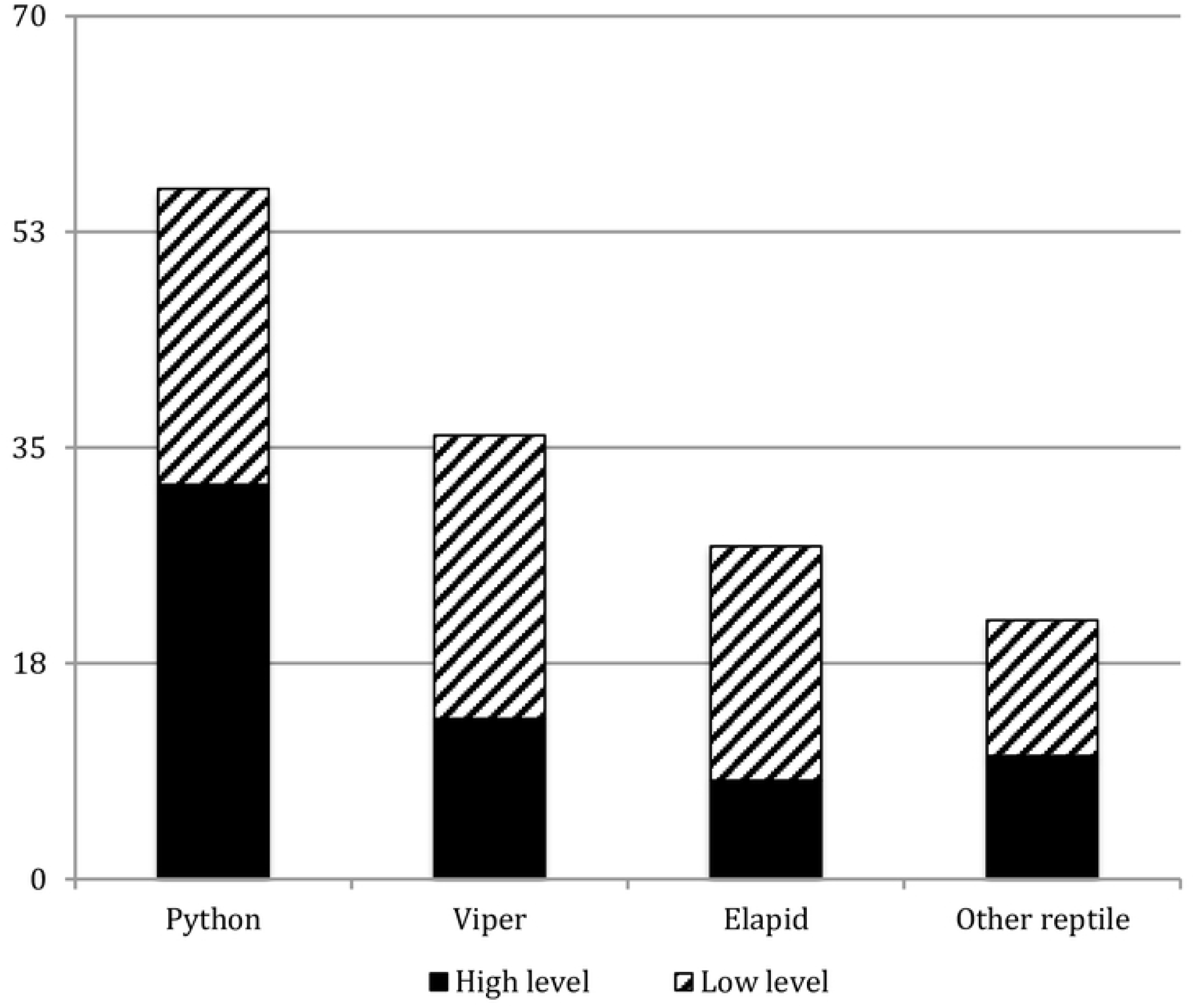
Frequency of interaction types with dangerous snakes and other reptiles identified to family at Fongoli, Senegal*. *High-level reactions include attack, charge, avoid/leave and vocalize at reptile while low-level reactions include ignoring or vigilance only.

A binary logistic regression revealed significant differences in high versus low-intensity responses to dangerous snakes according to whether they were arboreal or terrestrial (*n*=147, X^2^=16.699, df=4, p=0.002). Analyses of high versus low intensity reactions by chimpanzee age-sex class and snake type revealed significant differences according to age-sex class (*n*=123, X^2^=9.217, df=4, p=0.056). No other variables (snake size, snake activity) significantly affected chimpanzee reaction intensity.

A total of 35 different individual chimpanzees were observed to react to large reptiles at Fongoli (S1 Table 2). Apes appeared to vary in their reaction to reptiles, although small sample sizes precluded more detailed statistical analyses. Qualitatively, certain chimpanzees seemed to react more strongly than others towards snakes and even reptiles in general, with one adolescent male (DW) almost always using weapons against them. Certain adolescent males (DW, JM) consistently exhibited aggressive behavior towards reptiles. Of the individuals that interacted most with reptiles, four of the top five chimpanzees were adolescent males. However, an adult male (KL) was observed to encounter and/or interact with reptiles more than any other individual. Overall, 19 different males and 16 different females were seen to encounter and/or interact with reptiles out of a possible total of 47 different individuals within the study group between 2005 and 2015.

When qualitative comparisons among the three most commonly encountered dangerous snake types are made (vipers, elapids and pythons), some differences emerge (S1). The large elapids tend to elicit an avoidance response on the part of the chimpanzees involved in the interaction. Among the reactions commonly exhibited by chimpanzees that find themselves in close proximity to a cobra or mamba are “jumps back” or simply “leaves area” (encounters # 2, 5, 34, 44). Large vipers (those represented in the chimp encounters are mostly puff adders, *Bitis arietans*) often elicited an approach and view response with several instances of multiple chimps approaching the vicinity of the discovered snake, visually scanning the area until they were able to locate the immobile snake, and then leaving (encounters # 3, 10, 15, 24, 37, 45, 48). Pythons elicited the strongest response on the part of the chimps with individual encounters eliciting multiple attacks, alarm vocalizations and display behaviors on the part of the chimpanzees (encounters # 4, 6, 18, 24, 25, 27, 28, 31, 40, 41, 47, 50, 52).

In chimpanzee responses to elapid snakes, there are simply few observations recorded. Chimpanzees vocalize, view the snake, and then leave the area. There are incidents of aggression such as shaking objects at or attacking with objects, but these are limited. Based on a review of these encounters, it is clear that the reduced response is at least in part due to the brevity of the encounter. Mobile elapids turn up in close proximity to chimpanzees and when the chimpanzees respond to them, either the snake leaves the area, the chimpanzee leaves the area, or both of these things occur. In general, the main chimpanzee response is one of avoidance.

When the chimpanzees notice a viperid, and all of the recorded encounters (S1) are with the puff adder, *Bitis arietans,* a large bodied viper that combines strong crypsis with immobility, the response is somewhat different. Here, the chimpanzees often respond to discovery of the snake by approaching the vicinity of the snake and viewing the area where the snake is indicated by the first chimpanzee to call out its presence. They seem to be searching for the snake visually and, when found, they then often leave the area soon afterward. In some cases, the snake is approached quite closely, to within one meter, for a close view, and then left alone. Although vipers are occasionally struck at with objects, the chimpanzees give at best a mild fear response with little evidence of overt alarm or heightened anxiety.

The response to pythons is very different. Although there are two species of pythons in the Fongoli region, the recorded encounters are probably all, except one, with the rock python, *Python sebae,* which is clearly a potential predator of chimpanzees. In these encounters, the chimpanzees not only attack the snake more frequently than is seen with the other two types of snakes, but a variety of behaviors are reported in these encounters that are not reported with the other snakes and these are all behaviors that indicate increased levels of anxiety or aggressiveness on the part of the chimpanzees. These behaviors include approaching the snake while vocalizing, either with waa alarms or pant hoots (4, 25, 27, 50), males pant-hooting in chorus (4), slapping the substrate (4, 28), approaching the snake while piloerect (25, 28), chimpanzees chasing each other, and several types of displays, including with objects, while vocalizing, and while moving past the snake.

### Adult female chimpanzee dies of apparent snakebite

On 2 November 2012, research assistants found adult (estimated age, 17 years) female chimpanzee Tia (TI) lying supine on the ground about 10 meters away from her 2-month-old male infant Toto (TO). Her 4.5-year-old juvenile daughter Aimee (AM) was in trees nearby. Tia was lying supine; her right arm was extended, and she was holding onto the trunk of a small tree with her right hand. Her left arm was also extended, and a small tree was in the crook of her neck on the left side. Her pelvis was flat on the ground, legs extended, with the left one slightly bent at the knee. It appeared as though TI died during the night or early morning. Her abdominal region was slightly bloated, but there was no smell of decomposition. No scavengers had visited her body. There was insect activity around her eyes, including adult blowflies and perhaps egg masses. In humans, flies begin to colonize a corpse within minutes or hours after death [35]. At 1348 hours researchers approached TI, whose stomach was then more bloated, indicating putrefaction was occurring. Her left leg also looked somewhat swollen. There were four fresh wounds detected on her left foot; two were on the dorsal surface; they looked like puncture marks and were about 3 cm apart from each other (Fig. 5). The other two wounds looked like shallow scrapes, one each on the medial and lateral aspects of her foot. Tia was buried, and researchers retrieved TO for care. Additionally, TCL, Harry W. Greene and a veterinarian (M. Schaer) experienced with snakebites in domestic animals all concurred that TI likely died of a lethal snakebite. The placement of the punctures and lack of extensive edema and necrosis are indicative of a cobra or mamba bite rather than a viper bite. Given the distance between the punctures, this snake would have been quite large [36].

**Figure 5.**
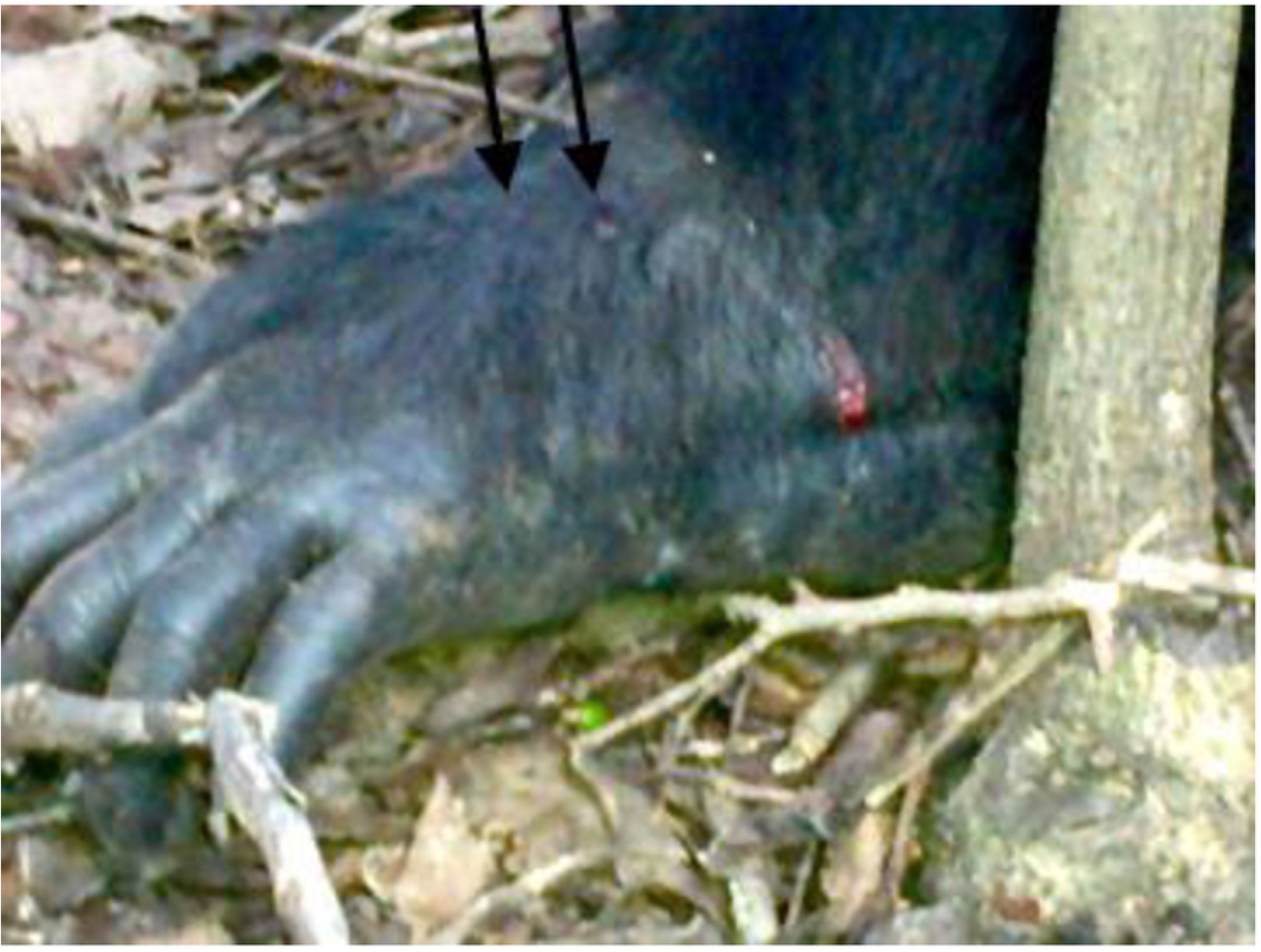
Photo of deceased adult female chimpanzee Tia’s foot*. *Arrows indicate 2 puncture wounds. Wound visible on lateral side of foot is a shallow scrape, likely made as foot was pulled through a narrow opening between boulders or tree branches.

## Discussion

Chimpanzees at Fongoli encounter snakes relatively frequently. Compared to Mahale chimpanzees in Tanzanian forest and woodlands, who reportedly encountered pythons only once in 40 years [37], Fongoli chimpanzees encountered pythons 14 times over the course of 10.25 years of study of habituated chimpanzee subjects. Bossou chimpanzees in Guinea, at a more forested site, were reported to encounter snakes only six times during almost 10,000 hours of observer contact with apes in 10 years [34], while Fongoli chimpanzees encountered snakes 37 times in 10.25 years. Researchers at Budongo, Uganda report encountering vipers approximately every two months, but do not report chimpanzee encounters with snakes; when presented with a snake model, these chimpanzees emitted alarm calls [38]. Additionally, almost all of the snakes that we observed chimpanzees to encounter in this study were dangerous (i.e., constricting or venomous). Compared to the frequency with which researchers encountered dangerous versus non-venomous snakes (see Table 1), this indicates perhaps that non-venomous snakes were ignored by Fongoli chimpanzees except in rare instances. In several cases an observer detected a snake while nearby apes apparently did not, although this was rare. In one case, an adult male seemed only to react to the (non-venomous) snake after the observer approached it (see S1, Table 1, case #35). Dangerous snakes, therefore, appear to be of concern to Fongoli chimpanzees, and they are not encountered infrequently. Detailed analyses of both snake type and chimpanzee age-sex class revealed further patterns regarding the potential selective pressures dangerous snakes present in a savanna-mosaic environment.

Our analyses of Fongoli chimpanzee reactions to snakes partly support the SDH. Based on multinomial logistic regression analyses, Fongoli chimpanzees did not vary in their reactions according to type of reptiles, but binary logistic regression analyses revealed that chimpanzee age-sex class and location of the snake did significantly affect high versus low intensity reactions. Various age-sex classes of apes reacted differently, with adolescent males being more aggressive, and all apes reacting more strongly to arboreal snakes compared to terrestrial ones. Other variables such as size and activity of the snake did not affect chimpanzee reaction intensity.

As Isbell [1, 2] notes, few accounts of predatory events by snakes on primates exist, with more sightings of constrictors preying on primates than deadly interactions between primates and venomous snakes (Table 1). However, the fact that snakes elicit predator-specific alarm calls from many primate species suggests this type of animal is sufficiently dangerous to evoke the evolution of acoustically distinct warning vocalizations [1]. The recent finding that snakes elicit significant responses in particular areas of the brain of Japanese macaques (*Macaca fuscata*) also supports Isbell’s [1] hypothesis that snakes were an important selective pressure acting on the evolution of the visual system and fear module of primates [39]. In fact, pythons still present a significant threat to humans living more traditional hunter-gatherer lifestyles even today [40]. Similarly, a survey of snakes in the Bandafassi region of Senegal, where the current study was conducted also indicates that snakes pose a threat to humans living here, causing 0.9% of all mortality but 28% of accidental deaths over a 24-year period [41]. One-third of the 34 snake species recorded in the Bandafassi study were dangerous [41]. Humans in this area commonly kill snakes they encounter, and instances of lethal bites received by humans are reported yearly, often in conjunction with wild plant food gathering season when people climb into vine-covered trees to collect fruits of the liana *Saba senegalensis*. A study of animal contact and injury around Kibale National Park, Uganda found that approximately 11% (*n*=20 of 181 injuries reported) of cases where animals caused injury to people living in and around forest fragments adjacent to the park involved snakes [42]. Clearly, in many areas of the world, even today snakes pose a significant risk to humans as well as other primates.

The fearful and defensive reaction of Fongoli chimpanzees directed towards snakes is not surprising given that during our relatively short study (i.e., less than one chimpanzee generation in length) one adult female (TI) appeared to have died from snakebite. Given that her 2-month-old infant (TO) was very unlikely to be adopted by a lactating female, researchers rescued him. He and his mother, as well as his juvenile sister (AM), who disappeared six months later, represent three of 15 of the deaths or disappearances recorded at Fongoli over the course of the eight years since all individuals were positively identified. These three cases of individuals removed from the gene pool can be attributed to snakebite, either directly (TI) or indirectly (TO, AM). Together they represent 20% of individuals lost from the Fongoli gene pool over the course of this study. Although the death/disappearance rate at Fongoli is low, the effects of a lethal snakebite can have a strong effect on a chimpanzee community. The significantly stronger reaction to arboreal snakes by Fongoli chimpanzees suggests even semi-terrestrial apes are at risk when arboreal.

Although the differences in the chimpanzees’ response to venomous versus constricting snakes do not differ statistically from one another, the collective response of the group to these three different types of snakes shows interesting qualitative differences. In particular, they respond to venomous snakes by either immediate avoidance, if the snake is an active species such as an elapid (cobra or mamba), or by first attempting to visually fix the position of the snake if it is a sedentary viper. This suggests that the chimpanzees recognize these latter type of snakes to be more of an environmental hazard that should be avoided than a serious threat. Even though individual Fongoli chimpanzees sometimes attacked such snakes, the response of the group is largely avoidance with mild expressions of fear or anxiety.

The main difference in the response to elapid snakes and vipers appears to be that the chimpanzees and elapids show a pattern of mutual avoidance that often results in relatively brief encounters that permit only a few behaviors of any kind to be recorded (notice the low numbers of behaviors recorded in Table S1), whereas viper encounters are often prolonged because the multiple chimpanzees will approach the area where the viper has been indicated, scan the area until the threat is visually fixed, and only then move away from it.

The response of the Fongoli chimpanzees to Rock Pythons, however, is clearly distinct. Even though these pythons are typically encountered when they are in a sedentary state, resting in a coil among dense branches and vines or in a water hole, the response of the chimpanzees typically indicates heightened anxiety and aggression, including piloerection and display behaviors that are similar to those by males when asserting dominance. Pythons are the only snakes that elicited several of the more aggressive types of responses by the chimpanzees, including approaching the snake while piloerect, slapping the substrate, approach while vocalizing, chorusing, and all types of display behavior (including display, display with object, display while vocalizing, and chimpanzees chasing each other) (S1).

Interestingly, our findings do not support the notion that non-human hominins consistently show a strong fear response to snakes. Rather than immediately avoiding snakes or fleeing from the vicinity when a group member gives a snake-specific warning call, the Fongoli chimpanzees very often were attracted to the site of a discovered snake. One of the most common responses of a party confronted with the presence of a snake was for each party member to approach the vicinity and cautiously search for the offending reptile. Once located visually, most of the chimpanzees would then leave the vicinity (aggressive responses that included attacking the snake with a branch or projectile also occurred, but were more limited). This suggests that the chimpanzees recognize snakes as a potential source of danger, but one that can be avoided once identified and located. There may be some parallels here with the chimpanzees’ response to fire (Pruetz and LaDuke, 2010). In the case of fire, chimpanzees were often seen feeding and behaving quite normally while in the path of an oncoming brush fire, leaving the area only when the fire approached to a very close proximity. Thus, the approach of a potentially dangerous entity whose properties the chimpanzees are familiar with does not generally elicit a strong fear response, but rather a response that is consistent with the need for a certain level of caution. In the same way, the chimpanzee’s response to snakes is clearly graded and proportional to the level of threat represented by the snake with the strongest response reserved for snakes that represent not just an environmental hazard, but are actually potential chimpanzee predators.

Although we agree with some aspects of Isbell’s SDH hypothesis, we find her focus on snakes as the primary force driving the evolution of the mammalian fear module to be overstated. Her reasoning for considering only crown clades of predators, thus excluding myriad potential extinct predatory forms as potential influences on the evolution of the mammalian fear module is unclear. The earliest mammals and their immediate ancestors clearly would have been influenced by predatory dinosaurs and the early avialians, whose feathers would have produced distinctive and recognizable patterns, not entirely unlike the scales of snakes. In fact, crocodilian and terrestrial dinosaurian predators would likely have been far more influential in the role of shaping the mammalian fear module. Thus, although snakes may have played an important role in the early evolution of the mammalian fear module, it is unlikely that they were the only group fulfilling this role.

Isbell’s carefully considered arguments regarding the evolution of the primate visual system in response to the potential threat of snake envenomation is sound and is supported by the findings of this study. In particular, encounters with venomous snakes, and especially the sedentary vipers, by the Fongoli chimpanzees often include bouts of visual scanning that involve many members of a group with individuals approaching the area where a snake has been seen by another, apparently scanning, or peering into the area until the snake is seen, and then moving on. That Fongoli chimpanzees seem to be compelled to search out and fix the position of a potentially dangerous object in their environment supports the notion that this knowledge has great survival value. In fact, locating the site where a snake is resting and then avoiding that area considerably reduces the threat that the snake poses because it is extremely unlikely that the snake will pursue an active chimpanzee.

## Conclusions

Our data on Fongoli chimpanzees’ natural reactions to potentially lethal reptiles provides insight into the degree to which snakes of different types, sizes, and locations are considered threats and could therefore reflect the degree to which such snakes serve as selective pressures in the present and perhaps in the past. Our results indicate snakes are serious threats in a savanna environment, and chimpanzees’ reactions to dangerous snakes vary regarding location of the reptile as well as the age-sex class of the apes. We found no statistical evidence of differential reaction by chimpanzees to constricting versus venomous snakes with our current sample. Perhaps relative to the intrinsic threat of dangerous snakes, qualitative differences in chimpanzees’ reactions to the three most threatening types of snakes suggest a subtly graded response that recognizes different levels of threat from the three categories of dangerous snakes most frequently encountered. Dangerous arboreal snakes of any type were met with significantly escalated reactions compared to terrestrial snakes. Adolescent males were significantly more intensively reactionary than chimpanzees of other age-sex classes. Finally, the loss of an adult female via a lethal snakebite and the subsequent loss of her two offspring from the population indicate the significant effects venomous snakes can have on chimpanzees at Fongoli.

## Acknowledgements

Thanks to the Republic of Senegal and the Department du Eaux et Forets for permission to conduct research in Senegal. Stacy Lindshield provided assistance with manuscript preparation. Dr. Michael Schaer provided helpful information on the likelihood of snakebite causing death in the adult female, Tia. Research assistants who aided in data collection include Dondo Kante, Stacy Lindshield, Michel Sadiakho, Erin Wessling, Waly Camara and Fiona Stewart. Harry W. Greene and Gordon Burghardt provided helpful comments on an earlier version of this manuscript.

